# YX0798 Is a Highly Potent, Selective, and Orally Effective CDK9 Inhibitor for Treating Aggressive Lymphoma

**DOI:** 10.1101/2024.12.31.629756

**Authors:** Vivian Jiang, Yu Xue, Hong Kim, Qingsong Cai, Tianci Zhang, Lei Nie, Joseph McIntosh, Yang Liu, Haiying Chen, Jia Zhou, Michael Wang

**Author notes:** equal contribution (Vivian Jiang, Yu Xue). **Correspondence** Dr. Michael Wang, Department of Lymphoma and Myeloma, The University of Texas MD Anderson Cancer Center, 1515 Holcombe Blvd., Houston, TX 77030, USA;.; Dr. Jia Zhou, Department of Pharmacology and Toxicology, University of Texas Medical Branch, Galveston, TX 77555, USA;.; Dr. Vivian Jiang, Department of Lymphoma and Myeloma, The University of Texas MD Anderson Cancer Center, 1515 Holcombe Blvd., Houston, TX 77030, USA;.

## Abstract

Non-genetic transcription evolution has been increasingly explored and recognized to drive tumor cell progression and therapeutic resistance. As the regulation hub of transcription machinery, cyclin-dependent kinase 9 (CDK9) is the gatekeeper of RNA polymerase II (Pol II) transcription, and CDK9 dysfunction results in transcriptomic reprogramming and tumor cell progression. We recently reported that the HSP90-MYC-CDK9 network drives therapeutic resistance in mantle cell lymphoma (MCL) through transcriptomic reprogramming. We also showed that targeting CDK9 by AZD4573 and enitociclib is a safe and effective treatment in preclinical MCL models, supporting CDK9 as a valid therapeutic target for MCL. However, current CDK9 inhibitors (CDK9i) under therapeutic development have room for improvement due to limited target selectivity and oral bioavailability. To this end, YX0798 was discovered to be a novel CDK9i through structural optimization. YX0798 demonstrated remarkable target selectivity and high affinity in binding to CDK9. Furthermore, YX0798 showed good oral bioavailability. YX0798, when administrated orally (5 mg/kg, daily), led to an efficacious anti-tumor activity *in vivo* and showed the potency in overcoming therapeutic resistance. Mechanistically, YX0798 downregulated the short-lived oncoprotein c-MYC and pro-survival protein MCL-1 as a common mechanism of CDK9 inhibition. Moreover, YX0798 disrupted the cell cycle and resulted in transcriptomic reprogramming, eventually leading to cell death. Together, these data demonstrate that YX0798 has oral bioavailability, exquisite selectivity, and anti-tumor potency that results from driving transcription reprogramming towards tumor cell killing.

**KEY POINTS:**

- YX0798 showed high selectivity, target binding affinity, and anti-tumor efficacy in overcoming resistance to various therapies.
- YX0798 had good oral pharmacokinetics and anti-tumor effects that resulted from transcription reprogramming towards tumor cell killing.

## Introduction

Non-genetic transcription reprogramming has been recognized to be a major driver of tumor progression and therapeutic resistance ^1-3^. Cyclin-dependent kinase 9 (CDK9) is a central regulator of RNA polymerase II (Pol II), which transcribes all protein-coding genes and many non-coding RNAs ^4^. CDK9 forms a heterodimer with cyclin T1 or T2 to form the positive transcription elongation factor b (P-TEFb), which phosphorylates Pol II to trigger the release of a paused Pol II ^5^. In addition, CDK9 mediates the initiation and termination stage of Pol II transcription ^6-8^. Because transcriptional dysfunction results in transcriptomic reprogramming and promotes cancer development, targeting CDK9 has the potential to overcome therapeutic resistance ^9^. In fact, CDK9 has been recognized as a therapeutic target in treating cancer and a few CDK9i are under clinical investigation ^10^. Recently, for mantle cell lymphoma (MCL), an aggressive subtype of non-Hodgkin’s lymphoma, we and others have reported that transcriptomic reprogramming is strongly associated with disease progression and therapeutic resistance ^11-13^. With MCL patients, next-generation sequencing showed that CDK9 expression was upregulated and the cancer hallmark MYC_TARGETS was highly enriched in individuals who relapsed post chimeric antigen receptor (CAR)-T-cell therapy compared with individuals who were CAR T-cell therapy naïve, suggesting that targeting CDK9 could overcome that resistance ^14^.

As CDK9 has long been recognized as a therapeutic target, small molecules have been generated to inhibit CDK9 but many have failed in clinical trials due to a lack of selectivity for cancer cells and indiscriminate binding with other CDKs ^10^. To improve this, more selective CDK9i AZD4573 ^15^ and enitociclib ^16,17^ have been developed and are effective in hematologic malignancy models, and specifically in MCL preclinical models by our group ^14,18^. These two CDK9i have advanced to Phase I/II clinical investigation, and early clinical results showed that CDK9 is a promising therapeutic target. However, both CDK9i are administrated by intravenous infusion and no oral bioavailability has been reported. Here, to develop a CDK9i with higher selectivity and oral bioavailability, we performed rational drug design, target profiling, anti-tumor evaluation, and oral bioavailability assessment. Among our candidates, YX0798 was identified as a potent, selective, and safe CDK9i with oral bioavailability that could be beneficial for treating cancer and improving patient outcomes.

## Methods

### Chemicals

YX0798 was designed and synthesized through computer-assisted structure-based drug design. Other compounds, enitociclib and AZD4573, were purchased from SelleckChem. For *in vitro* studies, the compounds were prepared in DMSO. For *in vivo* studies, compounds were prepared in 2% DMSO, 30% PEG-400, and 5% Tween-80.

### 3D molecular docking

Molecular docking was performed with Small Molecule Drug Discovery Suite (Schrödinger Release 2022-2, Schrödinger, LLC, New York, NY, 2019) ^30,31^. The co-crystal of CDK9 with AZD4573 (PDB ID: 6Z45) was obtained from RCSB PDB. The selected structure was preprocessed and optimized with Schrödinger Protein Preparation Wizard. The 20 Å grid box was generated using Glide Grid and centered on AZD4573. Compound YX0798 was created with Maestro and prepared with LigPrep to generate a suitable 3-D conformation for docking. Ligand docking was performed with Glide using SP mode with default settings. The docked pose was incorporated into Maestro pose viewer to visualize the interaction on the binding site.

### Kinome profiling and binding affinity

Kinome profiling of YX0798 at 100 nM was performed using the scanMax assay against a panel of 468 kinases and relevant mutants (Eurofins). Kinase selectivity of YX0798 was visualized using TREEspot software (Eurofins). The binding affinity of YX0798 and AZD4573 to CDK9 was determined using the kdELECT assay (Eurofins).

### Cell lines

MCL cell lines JeKo-1, JeKo-R (with acquired resistance to ibrutinib ^32^), JeKo BTK KD (with intrinsic ibrutinib resistance due to BTK depletion ^32^), Mino, Mino-R (with acquired resistance to venetoclax ^33^), Maver-1, and Z138 were maintained in RPMI 1640 medium supplemented with 1% penicillin/streptomycin, 25 mM 4-(2-hydroxyethyl)-1-piperazineethanesulfonic acid (HEPES), and 10% FBS (all from Sigma-Aldrich), and cultured in a CO2 incubator at 37°C. Cell lines were authenticated by single-nucleotide polymorphism profiling.

### Patient samples

Patient samples were collected from peripheral blood or apheresis after obtaining written informed consent and approval from the Institutional Review Board at The University of Texas MD Anderson Cancer Center (MDACC). The peripheral blood mononuclear cells (PBMCs) from healthy donors were purchased from Gulf Coast.

### Cell viability, cell cycle and apoptosis assays

Assays were performed as described previously^29^. Briefly, for the cell viability assay, cells were seeded at 10,000 cells per well in 96-well plates and exposed to either vehicle or compounds for 72 h. Cell lysis was conducted using the CellTiter-Glo Luminescent Cell Viability Assay Reagent, and luminescence was quantified employing the BioTek Synergy HTX Multi-mode microplate reader. For the cell cycle assay, MCL cells treated with CDK9i were stained with propidium iodide, and then subjected to flow cytometric analysis using the NovoCyte Flow Cytometer to determine the percentage of each cell cycle stage. For the apoptosis assay, MCL cells were treated with either vehicle or test compounds, stained with Annexin-V and propidium iodide, and then subjected to flow cytometric analysis to determine the percentages of Annexin-V positive cells. Data analysis was conducted using NoVo Express or FlowJo10. Each experiment was repeated at least three times.

### Measurement of cellular oxidative stress

Cellular reactive oxygen species (ROS) levels were measured using CellROX^TM^ Green Reagent (Invitrogen, C10444) according to the manufacturer’s manual. The cells treated with CDK9i were harvested and stained with CellROX^TM^ Green Reagent (5 µM). Flow cytometry analysis was performed to determine the intensity of cellular oxidative stress using the NovoCyte Flow Cytometer.

### Western blotting

These assays were performed as described previously^29^ Briefly, MCL cell lines were treated with vehicle, AZD4573 or YX0798 prior to harvest. Cells were then lysed on ice in lysis buffer for 30 min, then centrifuged at 4 °C for 10 min at 14,000 g; 4 × loading buffer was added to the lysates after protein concentrations were measured. Antibodies used to visualize proteins are shown in Supplementary Table S1. ImageJ was used to determine the band intensity for each protein.

### Efficacy evaluation in patient-derived organoid (PDO) models

This assay was performed as described previously ^34,35^. Briefly, the PDO models were established using primary samples collected from patients with MCL. To examine the drug sensitivity of primary MCL cells, the PDO models were treated with BTK inhibitors or CDK9i for 72 h. The cell viability was measured using CellTiter-Glo 3D assay kits (Promega, G9681) following the manufacturer’s protocol.

### In vivo efficacy assessment in patient-derived-xenograft models

These assays were performed as described previously ^29^. The Institutional Animal Care and Use Committee of The University of Texas MD Anderson Cancer Center approved the experimental protocols involving animals. Briefly, patient-derived xenograft (PDX) mouse models were established by intravenously inoculating MCL cells into NSG mice. PDX cells isolated from earlier generations were subcutaneously injected into NSG mice. When the tumors became palpable, the mice (n = 5 for each group) were treated with compounds as outlined in **Figure 4**. Tumor volumes, survival duration, and body weight were monitored. Tumor diameters were measured biweekly starting on Day 0 of tumor cell inoculation until the maximal tumor diameter reached 20 mm, and tumor volumes were calculated as V = (L × W × W)/2. The Institutional Animal Care and Use Committee of MDACC approved the experimental protocols.

### Metabolic and PK profile studies of YX0798

The liver microsome stability assay, the MDCK-MDR1 permeability assay, kinetics solubility determination, and the *in vivo* pharmacokinetics study of YX0798 were performed by a CRO service (BioDuro-Sundia, Irvine, CA).

### Bulk-RNA sequencing

JeKo-1 and Mino cells treated with vehicle or CDK9i (AZD4573 or YX0798) at the indicated concentration for 6 or 24 hr were harvested and subjected to bulk RNA sequencing and analysis as described previously^14^.

### Statistical analysis

All statistical analyses were conducted using R software (version 4.0.3) and GraphPad Prism (version 9). Two-sided two-sample t-test was used to compare differences between two groups. Results were considered statistically significant for p < 0.05 (*); p < 0.01 (**); p < 0.001 (***); and p < 0.0001(****).

## Results

### YX0798 is a novel CDK9i with higher target selectivity than AZD4573

To develop CDK9i candidates with high selectivity and oral bioavailability, we designed and generated more than 70 CDK9i (data not shown) based on CDK9i AZD4573 through multiple rounds of structural optimization. YX0798 (**Figure 1A**) was identified as the most potent CDK9i candidate based on anti-tumor efficacy screening *in vitro*. To visualize how YX0798 binds to CDK9, we performed YX0798-CDK9 molecular docking via Schrödinger using a AZD4573-CDK9 co-crystal structure (PDB: 6Z45). Similar to AZD4573, YX0798 binds to CDK9 in the ATP binding site. As shown in **Figure 1A**, YX0798 bound well in the CDK9 active site ^19^, including a hinge moiety and binding moiety deep in the pocket. The aminopyridine skeleton formed crucial dual hydrogen bonds with Cys106, which contributed to the high affinity of YX0798 with CDK9 (**Figure 1A right panel**). In addition, other key interactions, such as the hydrophobic interaction of the protein with the trifluoromethyl group, also contributes to the selective binding.

**Figure 1.**
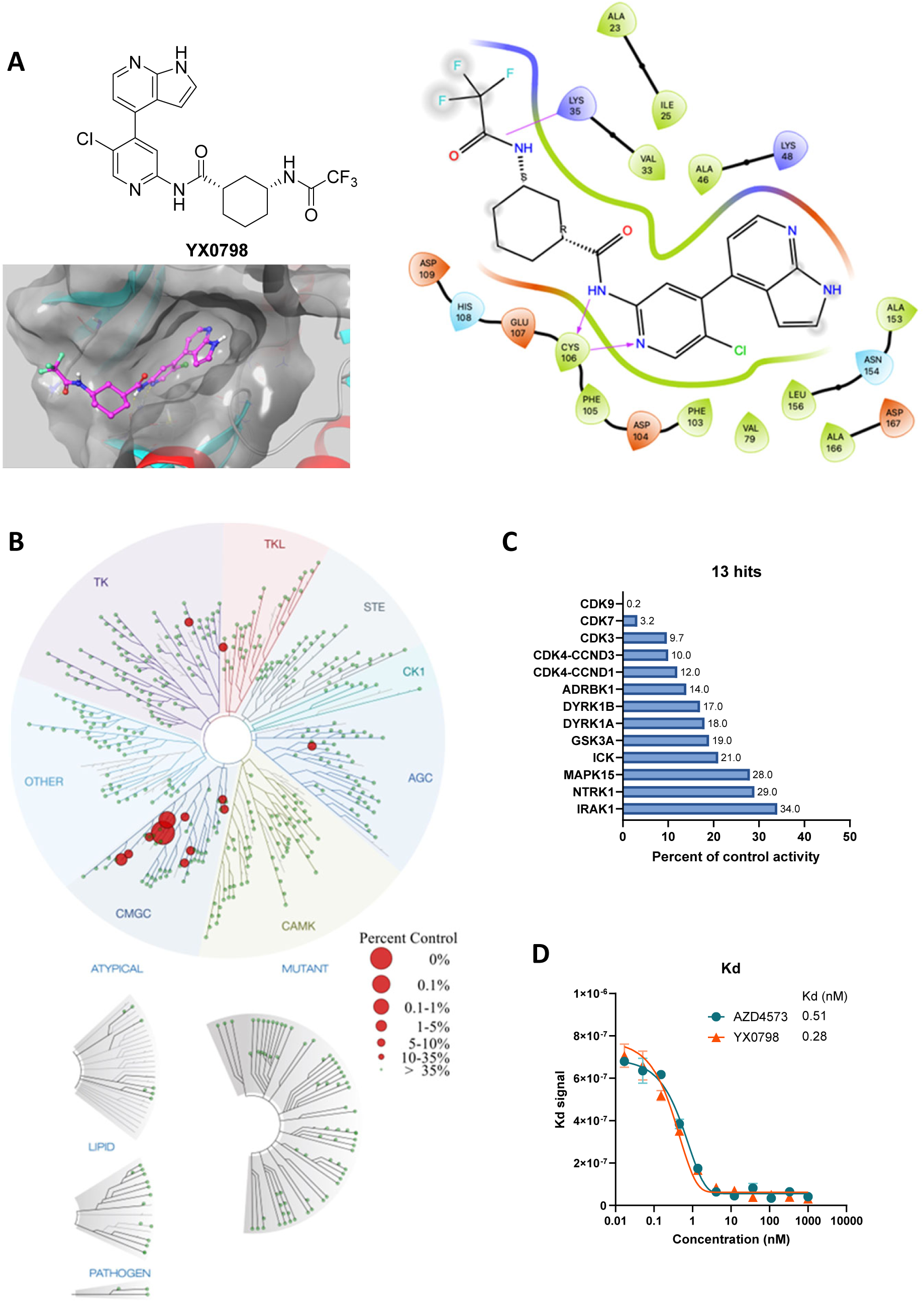
Identification of YX0798, a potent and selective CDK9i. (**A**) Chemical structure of YX0798 (upper left panel) and YX0798-CDK9 molecular docking. YX0798-CDK9 molecular docking via Schrödinger, using a publicly available AZD4573-CDK9 co-crystal structure (bottom left panel). YX0798 interaction with CDK9 visualized in the ATP binding pocket (right panel). (**B-C**) Kinome profiling of YX0798 at 100 nM using a scanMax panel consisting of 468 kinases (Eurofins) (**B**) and the thirteen hits identified with more than 66% kinase inhibition (**C**). (**D**) The binding affinity of CDK9i YX0798 and AZD4573 using a kdELECT assay (Eurofins).

To further determine the selectivity of YX0798, we performed kinome profiling using the scanMax assay, which includes a panel of 468 kinases. YX0798 at 100 nM led to only 13 hits with reduced activity ≥ 66% (**Figure 1B**). YX0798 led to a 99.8% reduction against CDK9 activity, which is 16-50-fold more selective for CDK9 than CDK7, CDK3, or CDK4 (**Figure 1C**). In contrast, AZD4573 was reported to have 16 hits with reduced binding ≥ 90% ^15^. Furthermore, YX0798 showed an extremely high binding affinity (*K*_d_ = 0.28 nM) (**Figure 1D**), which was 357-fold lower than the concentration used for the kinome profiling in **Figure 1C**, whereas AZD4573 showed a relatively weaker affinity (*K*_d_ = 0.51 nM) (**Figure 1D**). These results confirm that YX0798 has a greater CDK9 affinity and selectivity than AZD4573.

### YX0798 inhibited CDK9 activity and potently killed MCL cells *in vitro*

YX0798 (IC_50_ = 25.5-109.9 nM)) was highly potent in killing BTKi-sensitive MCL cells (Mino and JeKo-1) and BTKi-resistant cells (JeKo BTK KD and Z138) (**Figure 2A**), but not healthy PBMCs (**Figure 2B**). AZD4573 was more potent than YX0798 in killing MCL cells with a lower IC_50_ (6.5-13.8 nM) (**Supplementary Figure S1A**), which is not surprising considering the off-target effects of AZD4573 (**Figure 1B)**^15^. Indeed, we observed higher toxicity with AZD4573 in healthy PBMCs than YX0798 (**Supplementary Figure 1B**). Similar to AZD4573, YX0798 potently inhibited cell viability and induced apoptosis in all MCL cell lines assessed (**Figure 2C and Supplementary Figure S1C**). Western blots revealed that YX0798 inhibited CDK9 phosphorylation, induced poly-ADP ribose polymerase (PARP) and caspase 3 cleavage, and reduced anti-apoptotic MCL-1 in BTKi-resistant cell line Z138 (**Figure 2D-E**). YX0798’s potency was also demonstrated in BTKi-sensitive Mino and JeKo-1 cells (**Figure 2F**) and similar to that of AZD4573. Furthermore, reduction of phosphorylation of CDK9 substrate Pol II at Ser2, and non-CDK9 substrate AKT were seen in JeKo-1, Mino, and Z138 cells treated with YX0798 or AZD4573 (**Figure 2F**). In addition to MCL-1, c-MYC was reduced in a dose-dependent manner by YX0798 (**Figure 2F**), which we reported for AZD4573 and enitociclib in MCL cells ^15,16^ and in non-MCL hematopoietic cancer cells ^14,18^. Together, these data proved a common mechanism of action of CDK9i, leading to depletion of short-lived oncoprotein c-MYC and pro-survival MCL-1.

**Figure 2.**
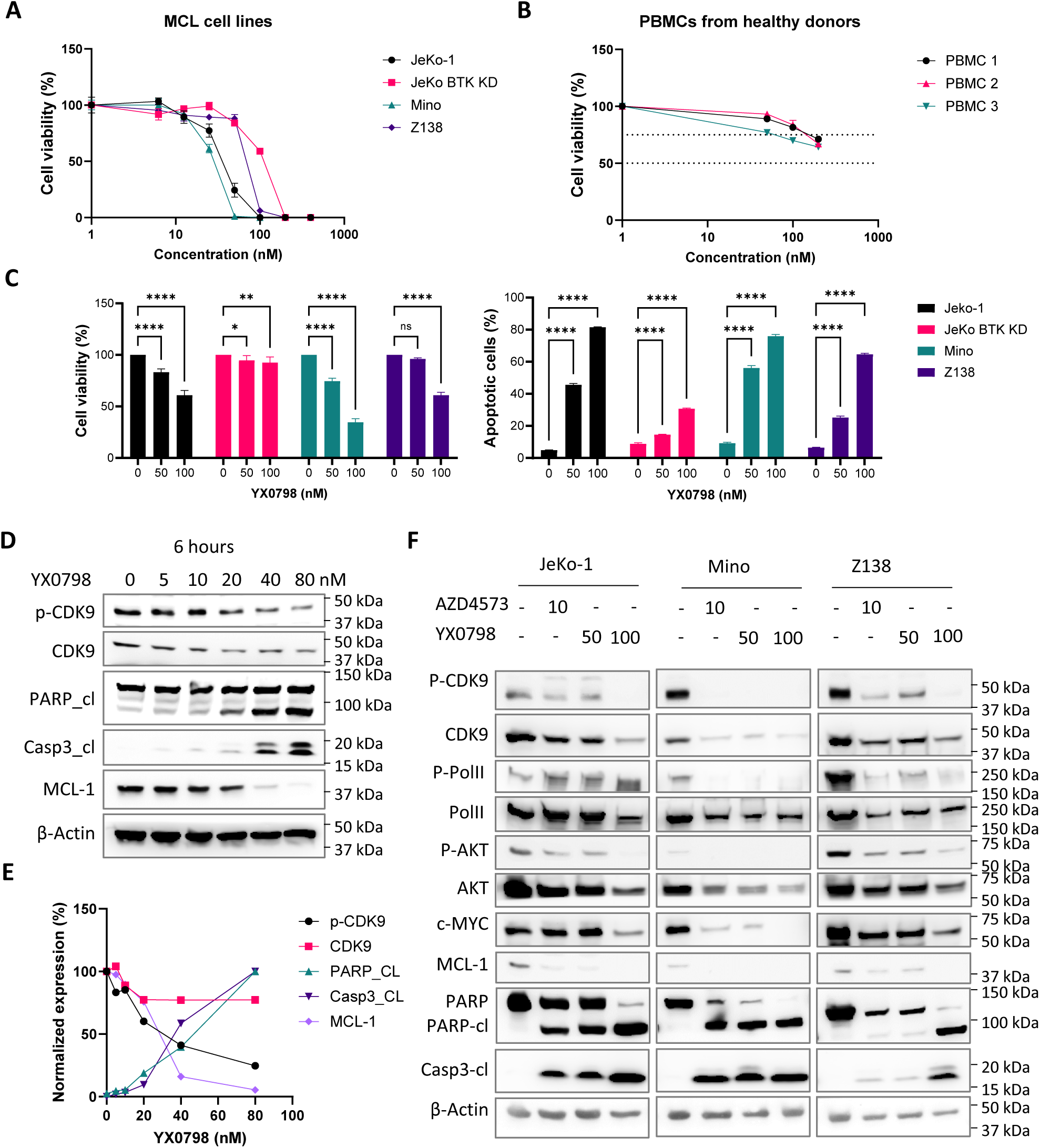
YX0798 inhibits CDK9 activity and is potent in killing MCL cells. (**A-B**) YX0798 potently inhibited cell viability of MCL cells (n = 4) (**A**), but not PBMC cells from healthy donors (n = 4) (**B**). (**C**) YX0798 inhibited cell viability and induced apoptosis in a dose-dependent manner. (**D-E**) YX0798 treatment for 6 hr inhibited CDK9 phosphorylation and induced cleavage of PARP and caspase 3, as well as MCL-1 depletion in a dose-dependent manner (**D**). The western band density was quantified by ImageJ and plotted (**E**). (**F**) The effects of YX0798 on protein profiling focusing on CDK9 pathways and apoptosis pathways. AZD4573 serves as a positive control. *, p < 0.05; **, p < 0.01; ***, p < 0.001; ****, p < 0.0001.

### YX0798 is stable, safe, and orally bioavailable

To further characterize the drug-like properties of YX0798, we performed *in vitro* liver microsome stability assays. YX0798 was stable in all three species tested with half-lives (T_1/2_ = 0.36, 0.58, and 0.94 h in mouse, rat, and human, respectively) (**Figure 3A**) comparable to AZD4573 (T_1/2_ = 0.18, 0.18, and 1.6 h, respectively) ^19^. YX0798 showed high cell permeability (P_app_ = 17.0 x 10^-6^ and 40.2 x 10^-6^ cm/s; efflux ratio = 2.4) as determined by the MDCK-MDR1 permeability assay (**Figure 3B**). YX0798 has high solubility (451 µM) in acidic FaSSGF (Fasted State Simulated Gastric fluid) buffer (pH1.2), much lower solubility (24.2 µM) in FaSSIF (Fasted State Simulated Intestinal fluid) (pH6.8), and no detectable solubility in pH7.4 buffer (**Figure 3C**). These data indicate that YX0798 is a basic drug given its pyridine core scaffold and high gastric fluid solubility. We next assessed the pharmacokinetic properties (PK) of YX0798 in rats following intravenous (iv) and oral administration. As shown in **Figure 3D**, YX0798 has a lower clearance rate (CL) and longer T_1/2_ than those of AZD4573 ^19^. Meanwhile, good plasma exposure after oral administration was seen, and the total drug exposure across time (AUC0-∞) values for YX0798 was 4457 ng·h/mL. YX0798 oral bioavailability (F%) is 16.5% and indicates the potential for oral administration, providing an advantage over other CDK9i such as AZD4573 and enitociclib with no oral bioavailability reported. The hERG inhibition assay, used to evaluate the potential cardiotoxicity of a drug candidate, showed that YX0798 has weak hERG inhibition at 10 µM (**Figure 3E**), which is much higher than the highest plasma concentration (2.61 µM) the compound can reach by oral administration in rats. Therefore, we do not expect that YX0798 will cause cardiotoxicity. Together, these data indicate that YX0798 is a stable and safe CDK9i with acceptable oral bioavailability.

**Figure 3.**
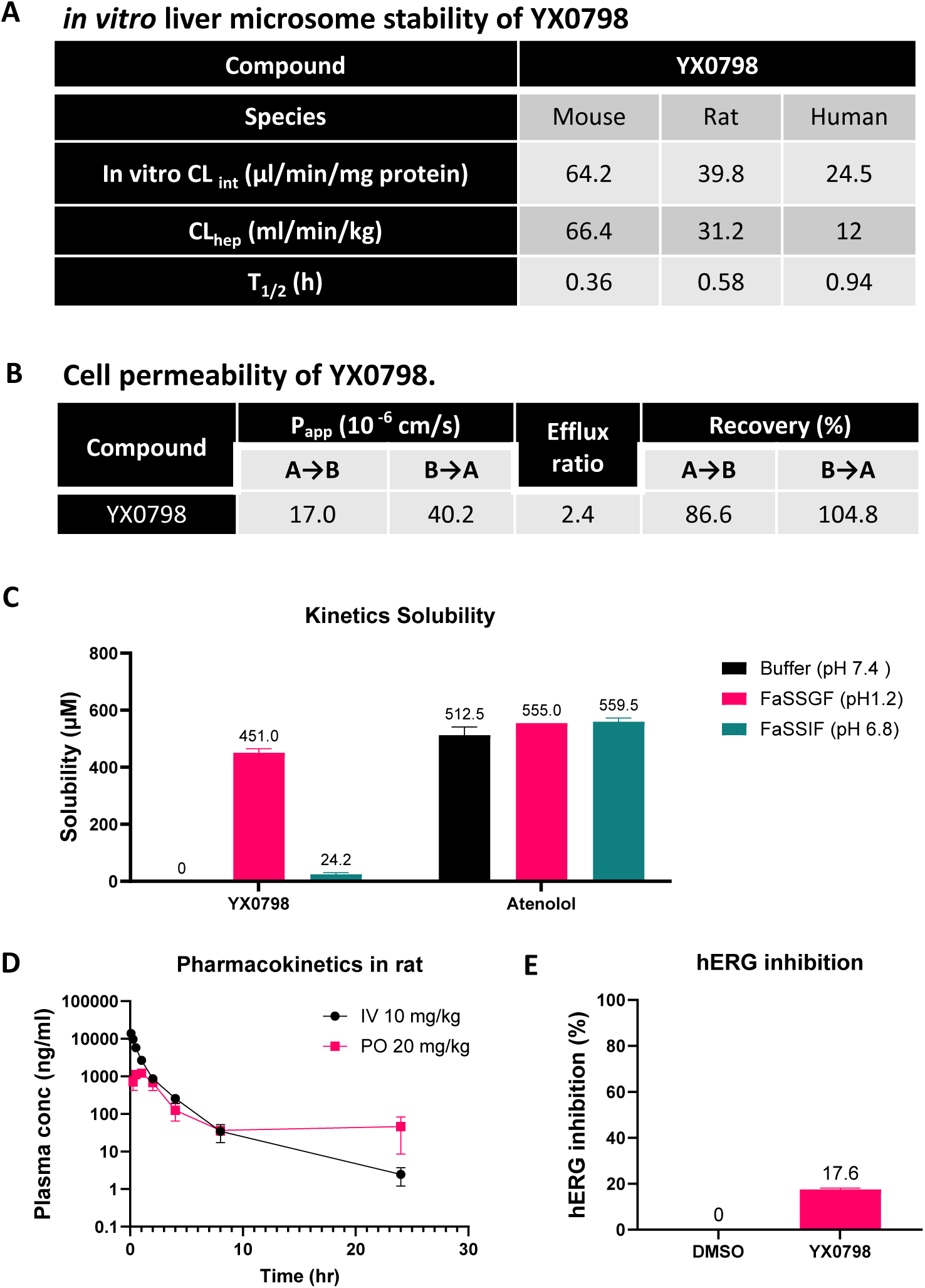
YX0798 shows oral bioavailability in rats. (**A**) YX0798 is metabolically stable in mouse, rat, and human as determined by an *in vitro* liver microsome stability assay. (**B**) YX0798 has high cell permeability. (**C**) YX0798 has good kinetic solubility in acidic FaSSGF (fasted state simulated gastric fluid). The beta-blocker atenolol was used as a control. (**D**) YX0798 showed oral bioavailability in rats. (**E**) YX0798 caused weak inhibition on hERG activity at 10 µM.

### Orally administrated YX0798 effectively inhibited the growth of patient MCL cells in cell culture and patient-derived models

To assess YX0798’s anti-MCL efficacy within a clinical context, we first assessed its efficacy in a sample from a patient who relapsed post pirtobrutinib. Consistent with the clinical response, the primary cultured cells no longer responded to BTK inhibitors ibrutinib, acalabrutinib, zanubrutinib, and pirtobrutinib, but was highly sensitive to CDK9i, using YX0798, AZD4573, or enitociclib (**Figure 4A-B**). As in MCL cell lines (**Figure 2D**), YX0798 induced dose-dependent inhibition of CDK9 phosphorylation, cleavage of PARP and caspase 3, and reduction of short-lived MCL-1 (**Figure 4C**) in the primary cultured cells. To further address the efficacy of YX0798, we established patient-derived organoid (PDO) models from a patient who relapsed post pirtobrutinib therapy. Consistent with the clinical response, these PDO models were resistant to BTKi, including pirtobrutinib and acalabrutinib. Interestingly, CDK9i AZD4573 and YX0798 were highly potent in killing these PDOs (**Figure 4D**).

**Figure 4.**
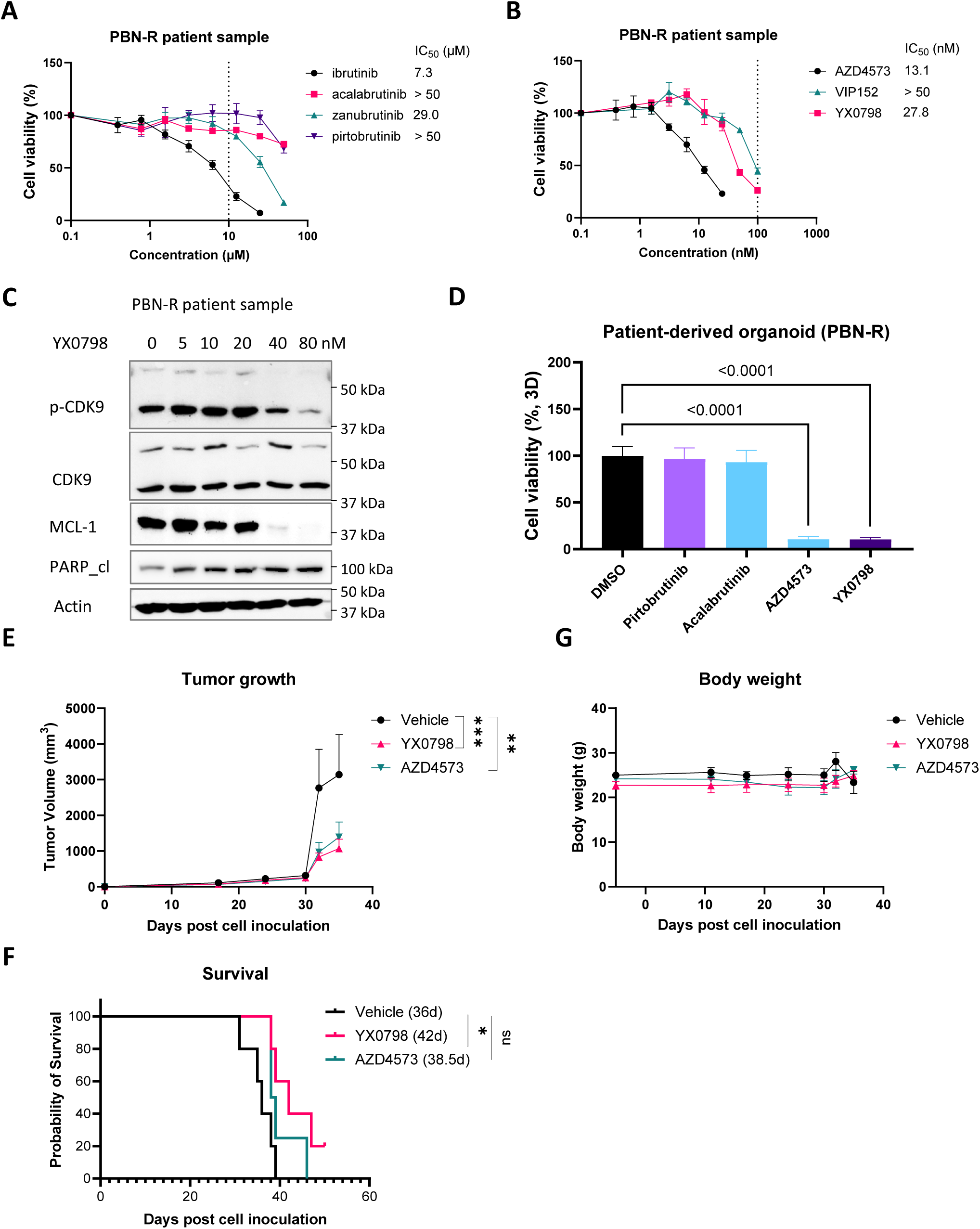
YX0798 is effective in inhibiting the tumor growth of PDX models in vivo. (**A-B**) One primary sample, originating from a MCL patient with relapse to pirtobrutinib, was confirmed to be resistant to BTKi, including ibrutinib, acalabrutinib, zanubrutinib, and pirtobrutinib (**A**), but was sensitive to CDK9i AZD4573, enitociclib, and YX0798. (**C**) YX0798 treatment for 6 hr inhibited CDK9 phosphorylation and induced cleavage of PARP, as well as MCL-1 depletion in a dose-dependent manner. (**D**) CDK9i AZD4573 or YX0798, but not BTK inhibition by pirtobrutinib or acalabrutinib, inhibited the growth of patient-derived organoid models, which originated from the same samples used in **A-C**. (**E-G**) Patient-derived xenograft models were established from a patient with sequential relapse to ibrutinib and CAR-T therapies. YX0798 was orally administrated at 5 mg/kg (5 days on and 2 days off every week, equivalent to 25 mg/kg/week), while AZD4573 was administrated intraperitoneally at 15+15 mg/kg (two doses at 2 hr apart, once a week, equivalent to 25 mg/kg/week). YX0798 treatment significantly inhibited the PDX growth (**E**) and prolonged mouse survival with better efficacy than AZD4573 (**F**). Treatment with AZD4573 or YX0798 did not affect mouse body weight (**G**). ns, not significant; *, p < 0.05; **, p < 0.01; ***, p < 0.001.

To assess YX0798’s anti-MCL efficacy *in vivo*, we generated a PDX model of relapsed MCL using a sample collected from a patient with sequential relapses to BTKi and CAR-T cell therapy. In the PDX models, YX0798 was orally administrated at 5 mg/kg (5 days on and 2 days off every week, equivalent to 25 mg/kg/week), while AZD4573 was administrated interperitoneally at 15+15 mg/kg (two doses at 2 hr apart, once a week, equivalent to 25 mg/kg/week). YX0798 showed marginally better efficacy in inhibiting tumor growth and prolonging mouse survival than AZD4573 (**Figure 4E-F)**. No apparent side effects were seen during any of the dosing regimens (**Figure 4G**).

### YX0798 disrupts cell cycling and metabolism

To understand the mechanism of action for YX0798, we performed bulk RNA sequencing. Mino and JeKo-1 cells were treated with YX0798 or AZD4573 for 6 hr and 24 hr (**Supplementary Figure S2A**). Consistent with the data in **Figure 2C**, YX0798 decreased cell viability in Mino and JeKo-1 cells in a dose-dependent manner as well as in a time-dependent manner (**Supplementary Figure S2B**). At 6 hr post treatment, YX0798 and AZD4573 started to inhibit cell viability, which became more prominent at 24 hr (**Supplementary Figure S2B**). For correlative studies, total RNA was isolated from aliquots of the same samples as in **Supplementary Figure S2A** and subjected to bulk RNA sequencing. We sought to identify the effects of AZD4573 or YX0798 on transcriptomic changes in MCL cells. Gene set enrichment analysis (GSEA) revealed the signaling pathways that were significantly downregulated or upregulated immediately at 6 hr in Mino and JeKo-1 cells, and expanded at 24 hr after CDK9 inhibition in JeKo-1 cells (**Figure 5A**). Multiple G2M-relevant and mitotic spindle-relevant pathways are the top downregulated pathways upon CDK9 inhibition, while cell metabolism-relevant pathways are the top upregulated pathways (**Figure 5A**). When JeKo-1 cells were treated with 50 nM YX0798 for 6 hr, multiple G2M-relevant and mitosis-relevant signaling pathways started to become downregulated, which became significantly more downregulated when treated with 100 nM of YX0798 for 6 hr (FDR < 0.05) (**Figure 5A-B and Supplementary Figure S2C**). Similar data were also seen in Mino cells (**Figure 5C and Supplementary Figure S2C**). These G2M signaling pathways were further downregulated when JeKo-1 cells were treated with 50 nM of YX0798 for 24 hr (**Figure 5A**). In addition, G2M pathways were downregulated upon AZD4573 treatment at 6 hr in Mino and JeKo-1 cells (**Figure 5A and Supplementary Figure S2C**). This cumulative downregulation was rarely seen with other downregulated pathways (**Figure 5A**). Together, these data indicate that downregulation of G2M signaling pathways is a predominant transcriptomic change with CDK9i AZD4573 or YX0798 in MCL cells, in addition to depletion of the oncoprotein c-MYC and pro-survival MCL-1 (**Figure 2D and 2F**). Because they are the primary pathways that were downregulated by CDK9i, they may drive the mechanism of action of CDK9i that leads to MCL cell death. For example, single-sample GSEA revealed that FISCHER_G2_M_CELL_CYCLE was significantly downregulated by CDK9i (**Figure 5B**). The inhibitors decreased the prevalence of cell cycle G2M pathways with enhanced effects when the concentration of YX0798 was increased or the treatment time was longer (**Figure 5B**). These data led to our hypothesis that CDK9i disrupts cell cycle progression and leads to cell death. We next checked whether CDK9i lead to cell cycle arrest. Indeed, treatment with either YX0798 or AZD4573 induced cell cycle arrest at the G0/G1 phase, not only in Mino and JeKo-1 cells but also in other more aggressive MCL cells, including BTKi-resistant JeKo BTK KD, Maver-1, and Z138 cells, as well as in Mino-R cells with acquired venetoclax resistance (**Figure 5C**).

**Figure 5.**
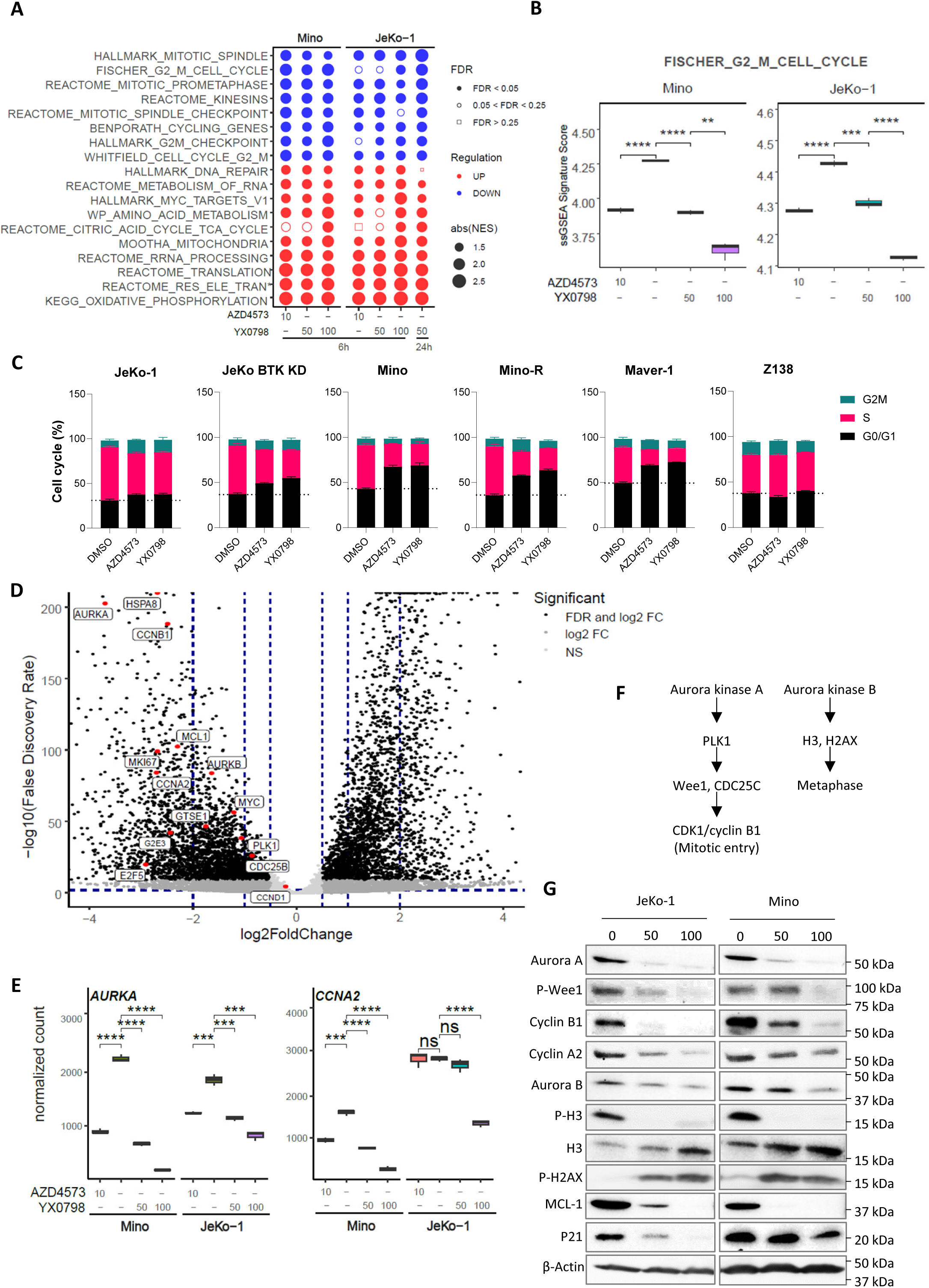
YX0798 disrupts cell cycling in MCL cells toward cell cycle arrest. (**A**) The bubble plots show top cancer hallmarks and signaling pathways downregulated (blue) or upregulated (red) upon CDK9 inhibition. (**B**) The FISCHER_G2_M_CELL_CYCLE pathway was significantly downregulated by YX0798 in JeKo-1 cells (50 and 100 nM for 6 and 24 hr) and Mino cells (50 and 100 nM for 6 hr). AZD4573 serves as a control. (**C**) YX0798 and AZD4573 induced cell cycle arrest in MCL cells as indicated. (**D**) Volcano plot shows the significantly altered genes upon YX0798 treatment at 100 nM for 6 hr. (**E**) Representative genes *AURKA* and *CCNA2* were significantly downregulated with CDK9i YX0798 or AZD4573. (**F**) Illustration of aurora kinase A and B mediated cell cycle progression. (**G**) Western blots confirmed the cell cycle disruption induced by treatment with YX0798 and AZD4573 for 6 hr. ns, not significant; *, p < 0.05; **, p < 0.01; ***, p < 0.001; ****, p < 0.0001.

To identify pivotal G2M genes responsible for cell cycle disruption by CDK9i, we performed differential gene expression analysis. Genes found to be significantly affected by YX0798 in MCL cells included *HSP8A* (Heat Shock Protein Family A (Hsp70) Member 8)*, AURKA* (Aurora Kinase A)*, CCNA2* (Cyclin A2), *CCNB1* (Cyclin B1), *CDC25B* (Cell Division Cycle 25B), *PLK1* (Polo Like Kinase 1), *E2F5* (E2F Transcription Factor 5), *MKI67* (Marker Of Proliferation Ki-67), *G2E3* (G2/M-Phase Specific E3 Ubiquitin Protein Ligase), and *GTSE1* (G2 And S-Phase Expressed 1) (**Figure 5D**). They were further validated by quantitative real-time PCR (**Supplementary Figure S3**). With CDK9i, *AURKA* and *CCNA2* mRNA decreased at 6 hr post treatment in Mino and JeKo-1 cells (**Figure 5E**). The AURKA-PLK1-Wee1-CDK1/CCNB1 axis is critical for mitotic entry and centrosome maturation, while AURKB-dependent histone H3 phosphorylation at Ser10 is the hallmark of mitosis ^20^ (**Figure 5F**). To further address the mechanism of action of CDK9i leading to cell cycle disruption, we checked G2M-relevant proteins with western blotting (**Figure 5G**). Protein expression of AURKA, CCNA2, CCNB1 and MCL-1 were reduced with CDK9i at 6 hr, accompanied by reduced phosphorylation of Wee1 and histone H3 (**Figure 5G**). Together, these data suggested that Aurora kinases may be the primary players responsible for downregulated G2M pathways. To address this, we assessed if inhibition of aurora kinases was effective for killing MCL cells. Based on a library screen of 320 FDA-approved/investigational agents, all four AURKA inhibitors effectively inhibited the viability of all MCL cell lines (**Supplementary Figure S4A**). Alisertib was especially effective in killing MCL cells (**Supplementary Figure S4B**). These data support the idea that targeting aurora kinases is an alternative approach for MCL treatment.

In addition to cell cycle disruption, CDK9i AZD4573 or YX0798 resulted in a remarkable enrichment of RESPIRATORY_ELECTRON_TRANSPORT and OXIDATIVE_PHOSPHORYLATION (OXPHOS) pathways, as well as other cell metabolism pathways (**Figure 5A and 6A**). Mitochondria are the main source of cellular ROS, which is produced by the electron transport chain during oxidative phosphorylation^21^. Therefore, it is likely that enrichment of these pathways leads to ROS overproduction that irreversibly damages cellular components, including lipids, proteins, and DNA, leading to apoptosis ^22,23^ (**Figure 6B**). Indeed, ssGSEA revealed that the REACTOME_ROS_AND_RNS_PRODUCTION_IN_PHAGOCYTES pathway was elevated by CDK9i YX0798 or AZD4573 (**Figure 6C**). Cellular ROS levels started to elevate at 6 hr and continued to climb at 24 hr (**Figure 6D**). These findings correlate well with reduced cell viability post treatment (**Figure 6D and Supplementary Figure S2B**). Moreover, H2AX phosphorylation, an early indicator of DNA damage ^24^, is increased with YX0798 (**Figure 5G**). These data demonstrate that CDK9 inhibition in MCL cells leads to disrupted cell cycling and metabolism, which may orchestrate eventual MCL cell killing.

**Figure 6.**
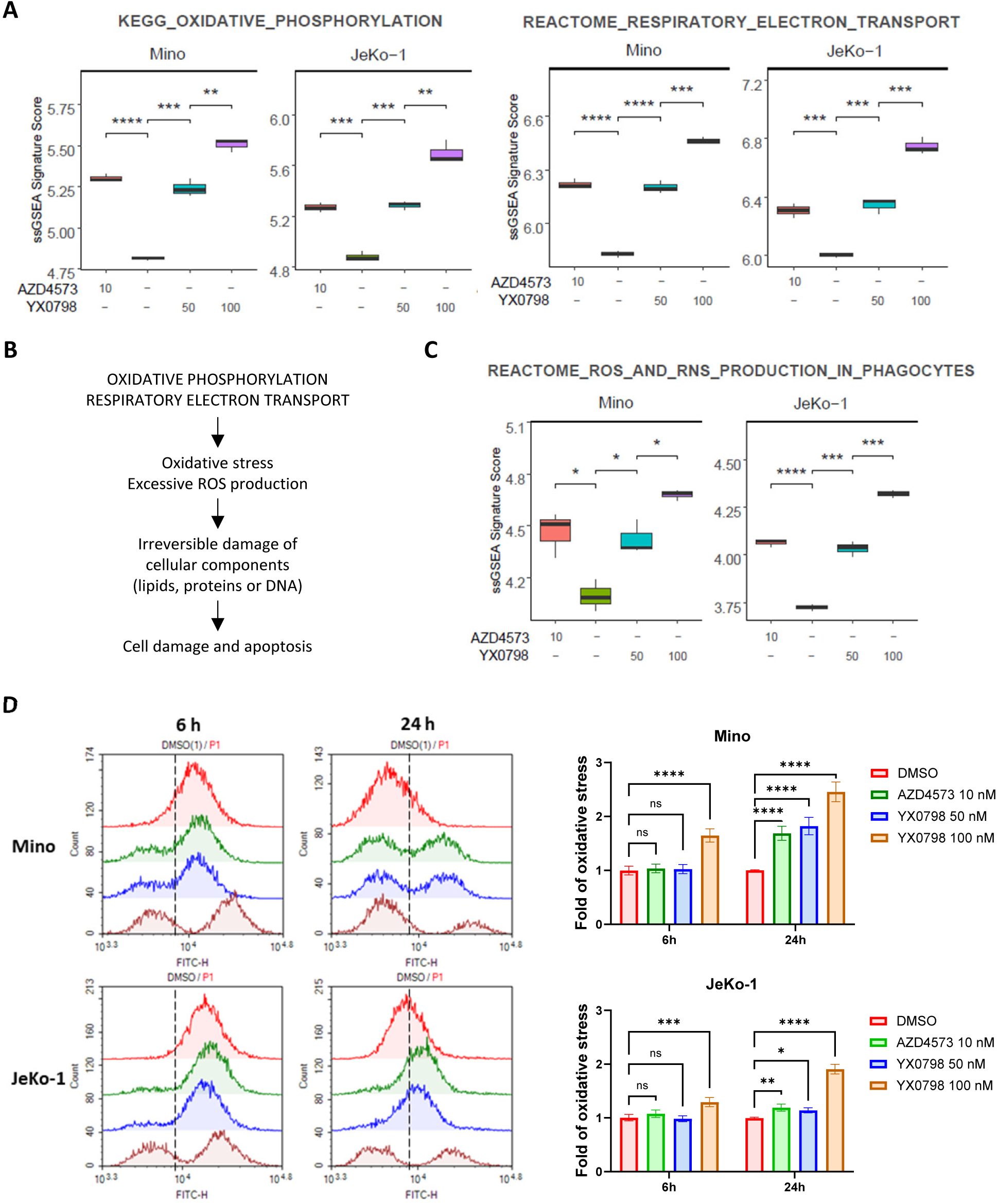
YX0798 induces enrichment of OXPHOS and excessive oxidative stress in MCL cells. (**A**) Enriched signaling pathways upon treatment with YX0798 and AZD4573 for 6 hr. (**B**) Illustration of enrichment of oxidative phosphorylation and RESPIRATORY_ELECTRON_TRANSPORT pathways leading to excessive oxidative stress and ROS production. (**C**) Enrichment of REACTOME_ROS_AND_RNS_PRODUCTION_IN_PHAGOCYTES signaling pathway upon treatment with YX0798 and AZD4573 for 6 hr. (**D**) ROS production is elevated upon treatment with YX0798 and AZD4573 for 6 and 24 hr in Mino and JeKo-1 cells. Left panels, the overlaid flow cytometry results of cellular ROS levels in cells. Right panels, the quantitative results of cellular ROS levels in cells. ns, not significant; *, p < 0.05; **, p < 0.01; ***, p < 0.001; ****, p < 0.0001.

## Discussion

CDK9 has long been recognized as a valuable target for therapeutic development in treating cancer ^10^. Early compounds that targeted CDK9 were essentially pan-CDKi, targeting multiple CDKs. For example, flavopiridol targets CDK1, CDK2, CDK4, and CDK6, and less potently targets CDK9 ^25,26^, whereas seliciclib targets CDK1, CDK2, and CDK5 and less potently targets CDK9 ^27^. Most pan-CDKi show suboptimal clinical benefits with adverse effects, and have therefore failed in clinical trials ^10^. The reasonable choice was to develop exquisite CDK9-specific inhibitors with limited off-target effects. The recent generation of CDK9i such as AZD4573 ^19^ and enitociclib ^16^ demonstrate high-CDK9 selectivity and are currently under clinical investigation; however, AZD4573 and enitociclib can only be administered by intravenous infusion. To overcome this limitation, we developed YX0798 as a potent and selective CDK9i with acceptable oral bioavailability. Compared to AZD4573, YX0798 showed higher selectivity in binding to CDK9. Furthermore, YX0798 demonstrated oral bioavailability that led to anti-tumor efficacy *in vivo* for overcoming therapeutic resistance. These results show YX0798’s advantages over other CDK9i including AZD4573 and enitociclib, with no reported oral bioavailability.

For AZD4573 and enitociclib, suppressing the expression of oncogenic c-MYC and pro-survival MCL-1 is the primary mechanism of action in hematologic malignancies ^16,19^. Consistent with these findings, dramatic protein depletion of c-MYC and MCL-1 was detected in MCL cells upon treatment with YX0798 (**Figure 2D-F**). We also observed differential changes in the mRNA expression of *MYC* and *MCL-1* upon treatment with YX0798 at different concentrations (**Supplementary Figure S2E, bottom two panels**). Furthermore, enrichment of the HALLMARK_MYC_TARGETS_V1 pathway is also seen with YX0798 or AZD4573 treatment in Mino and JeKo-1 cells (**Figure 5A**). This is somewhat unexpected given that the CDK9i mechanism of action leads to a genome-wide transcriptional pause, resulting in rapid turnover of short-lived c-MYC protein and mRNA ^16,19^. However, studies using other CDK9i, such as i-CDK9 and flavopiridol, reported an unexpected increase in c-MYC expression in other cancer cell lines due to a BRD4-dependent compensatory pathway ^28^. Our data further confirm that an increase of c-MYC expression is a common mechanism across CDK9i.

In addition to CDK9’s action on c-MYC and MCL, intriguingly, our transcriptomic profiling discovered that CDK9i YX0798 and AZD4573 resulted in remarkable downregulation of G2M signaling pathways, which likely leads to cell cycle arrest at the G0/G1 phase in MCL cells (**Figure 5B-C**). Cell cycle disruption has not been reported for earlier generations of CDK9-specific inhibitors such as AZD4573 and enitociclib. Therefore, this is the first time that cell cycle disruption has been revealed as a primary mechanism of CDK9-specific inhibition, not only for AZD4573 but also for our compound YX0798, indicating a common mechanism of CDK9i-induced tumor cell death.

Furthermore, our transcriptomic profiling revealed that oxidative phosphorylation and mitochondrial RESPIRATORY_ELECTRON_TRANSPORT pathways were enriched by CDK9i AZD4573 and YX0798 in Mino and JeKo-1 cells. Our recent translational studies using primary patient samples repeatedly revealed oxidative phosphorylation is associated with therapeutic resistance to BTKi and venetoclax ^13,29^. Therefore, it is intriguing that oxidative phosphorylation is a top enriched pathways upon CDK9 inhibition, which is commonly shared by YX0798 and AZD4573. Cellular ROS is predominantly produced by mitochondria, with ROS originating from the electron transport chain during OXPHOS process^21^. Therefore, it is not surprising to see that the REACTOME_ROS_AND_RNS_PRODUCTION_IN_PHAGOCYTES is also enriched upon CDK9 inhibition. Together, our findings may explain why we detected excessive cellular ROS upon CDK9 inhibition. Overproduction of ROS leads to damage of cellular components including DNA ^22,23^, as evident by H2AX phosphorylation (**Figure 5G**).

Collectively, our study demonstrated that YX0798 is a highly potent and selective CDK9i with oral bioavailability that not only depletes the expression of oncogenic c-MYC and pro-survival MCL-1, but also drives transcriptomic reprogramming towards disruption of cell cycle and metabolism (**Supplementary Figure S5**).

## Acknowledgements

This work is supported by a grant from the Cancer Prevention and Research Institute of Texas (CPRIT) (RP210062, MW and JZ)), an Institutional Research Grant (IRG, VJ) award from MD Anderson Cancer Center, the Steve and Nancy Fox Cancer Research Fund (MW), the John D. Stobo, M.D. Distinguished Chair Endowment (JZ), and the Edith & Robert Zinn Chair Endowment in Drug Discovery (JZ). We thank the patients and their families who contributed to this research study. We also thank Tracey L Baas for technical editing.

## Authorship Contributions

MW, JZ, and VJ conceived the idea. VJ, JZ, and XY designed the experiments; VJ, YX, HK, TJ, LN, JM, YL, and HC performed the experiments; VJ, YX, HK, and QC performed data analysis; VJ and XY directed and coordinated the project and wrote the manuscript; VJ, XY, JZ, and MW contributed to the manuscript revision. MW, JZ, and VJ acquired the funding. All authors read and approved the final manuscript.

