## Supplemental materials for "YX0798 Is a Highly Potent, Selective, and Orally Effective CDK9 Inhibitor for Treating Aggressive Lymphoma"

*, equal contribution

**This PDF file includes:**

- Materials and Methods
- Supplementary Tables S1-S2
- Supplementary Figures S1-S5
- References

### Materials and Methods

#### *cDNA synthesis and quantitative real-time PCR*

The method for cDNA synthesis and quantitative real-time PCR has been described elsewhere [1]. Briefly, first-strand cDNA was synthesized via the iScript cDNA Synthesis Kit (1708890, Bio-Rad) according to the manufacturer's manual. Quantitative real-time PCR was conducted using SsoAdvanced Universal SYBR Green Supermix (1725271, Bio-Rad) according to the manufacturer's manual. The primers used to amplify genes are shown in Supplementary Table S2. mRNA expression was normalized to *ACTB* and further normalized to one of the DMSO treated samples.

#### *High-throughput drug screen*

The method for the high-throughput drub screen has been previously described [2]. Briefly, a library of 320 FDA-approved/investigational drugs from Selleckchem (Houston, TX) was used for a high throughput drug screening. Four cell lines (JeKo-1, Mino, JeKo-R and Z138) were treated with each drug at 5 µM for 72 hr, and cell viability was determined using the CellTiter-Glo Luminescent Cell Viability Assay Reagent, and luminescence was quantified as described previously[2].

### Supplementary Tables

#### Supplementary Table S1: Antibodies used for western blots.

| Protein | Antibody | Company | Catalog # |
|---|---|---|---|
| P-CDK9 | Phospho-CDK9 (Thr186) | Cell Signaling Technology | 2549S |
| CDK9 | CDK9 (C12F7) | Cell Signaling Technology | 2316S |
| PARP-cl | Cleaved PARP (Asp214) (D64E10) | Cell Signaling Technology | 5625S |
| Casp3-cl | Cleaved Caspase-3 (Asp175) | Cell Signaling Technology | 9661S |
| MCL-1 | Mcl-1 Antibody | Cell Signaling Technology | 4572S |
| β-Actin | β-Actin (C4) | Santa Cruz | SC47778 |
| P-PolII | Phospho-Rpb1 CTD (Ser2/5) | Cell Signaling Technology | 4735S |
| PolII | Rpb1 CTD (4H8) | Cell Signaling Technology | 2629S |
| P-AKT | Phospho-Akt (Ser473) (D9E)XP | Cell Signaling Technology | 4060S |
| AKT | Akt (pan) (C67E7) | Cell Signaling Technology | 4691S |
| C-MYC | c-Myc Antibody | Cell Signaling Technology | 9402S |
| Aurora A | AURAurora A (D3E4Q) Rabbit mAb | Cell Signaling Technology | 14475S |
| Aurora B | Aurora B/AIM1 Antibody | Cell Signaling Technology | 3094S |
| P-Wee1 | Phospho-Wee1 (Ser642) (D47G5) Rabbit mAb | Cell Signaling Technology | 4910S |
| Cyclin B1 | Cyclin B1 Antibody | Cell Signaling Technology | 4138S |
| Cyclin A2 | Cyclin A2 (BF683) Mouse mAb | Cell Signaling Technology | 4656S |
| P-H3 | Phospho-Histone H3 (Ser10) (D2C8) XP® Rabbit mAb | Cell Signaling Technology | 3377S |
| H3 | Histone H3 (1B1B2) | Cell Signaling Technology | 14269S |
| p21 | p21 Waf1/Cip1 (12D1) | Cell Signaling Technology | 2947S |
| P-H2AX | Phospho-Histone H2A.X (Ser139) | Cell Signaling Technology | 2577S |

**Supplementary Table S2: primers used for qPCR.**

| Gene name | Forward primer | Reverse primer |
|---|---|---|
| CCNB1 | AATAAGGCGAAGATCAACATGGC | TTTGTTACCAATGTCCCCAAGAG |
| CCNA2 | CGCTGGCGGTACTGAAGTC | GAGGAACGGTGACATGCTCAT |
| CCND1 | GCTGCGAAGTGGAAACCATC | CCTCCTTCTGCACACATTTGAA |
| CDC25B | ACGCACCTATCCCTGTCTC | CTGGAAGCGTCTGATGGCAA |
| PLK1 | AAAGAGATCCCGGAGGTCCTA | GGCTGCGGTGAATGGATATTTC |
| AURKA | GAGGTCCAAAACGTGTTCTCG | ACAGGATGAGGTACACTGGTTG |
| AURKB | CAGTGGGACACCCGACATC | GTACACGTTTCCAAACTTGCC |
| E2F5 | ATGTCTTCTGACGTGTTTCCTC | CGGGGTAGGAGAAAGCCTT |
| MKI67 | ACGCCTGGTTACTATCAAAAGG | CAGACCCATTTACTTGTGTTGGA |
| G2E3 | CTGGTGACTCACAGAACCTTG | TGCCTCTCTGCCAAATTCCAC |
| GTSE1 | CAGGGGACGTGAACATGGATG | ATGTCCAAAGGGTCCGAAGAA |
| BIRC5 | AGGACCACCGCATCTCTACAT | AAGTCTGGCTCGTTCTCAGTG |
| Actin | CGGAACCGCTCATTGCC | ACCCACACTGTGCCCATCTA |

**Figures**

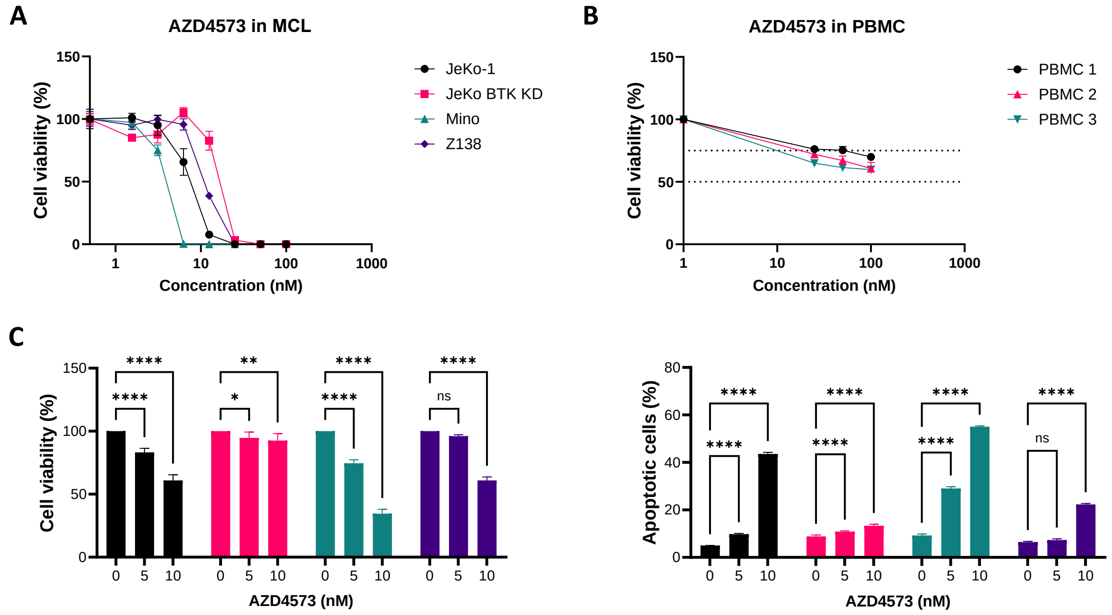


**Figure S1. The anti-tumor efficacy and toxicity of AZD4573.**

(**A-B**) The *in vitro* efficacy (**A**) and toxicity (**B**) of AZD4573 in MCL cell lines (n = 4) and in PBMC cells from healthy donors (n = 3). (**C**) AZD4573 inhibited cell viability and induced apoptosis in a dose-dependent manner.

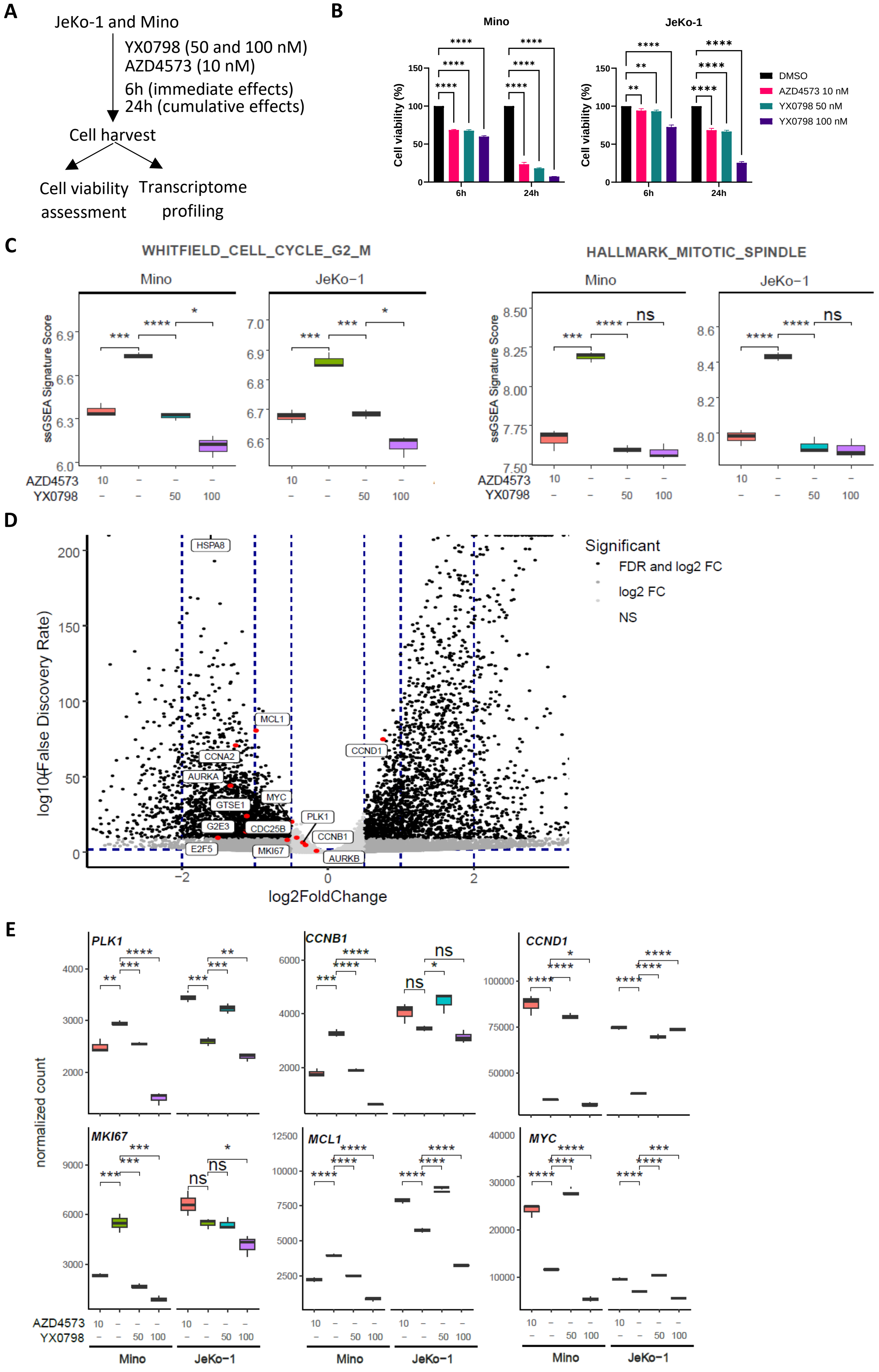
A
JeKo-1 and Mino
YX0798 (50 and 100 nM)
AZD4573 (10 nM)
6h (immediate effects)
24h (cumulative effects)
Cell harvest
Cell viability assessment
Transcriptome profiling
B
Mino
JeKo-1
Cell viability (%)
6h
24h
DMSO
AZD4573 10 nM
YX0798 50 nM
YX0798 100 nM
C
WHITFIELD_CELL_CYCLE_G2_M
HALLMARK_MITOTIC_SPINDLE
Mino
JeKo−1
ssGSEA Signature Score
AZD4573
YX0798
D
Significant
FDR and log2 FC
log2 FC
NS
log10(False Discovery Rate)
log2FoldChange
HSPA8
MCL1
CCND1
CCNA2
AURKA
GTSE1
MYC
PLK1
G2E3
CDC25B
CCNB1
E2F5
MKI67
AURKB
E
PLK1
CCNB1
CCND1
MKI67
MCL1
MYC
normalized count
Mino
JeKo−1

**Figure S2. CDK9i YX0798 downregulates G2/M-relevant signaling pathways.** (**A**) Illustration of experiment design. Mino and JeKo-1 cells treated with YX0798 (50 and 100 nM) or AZD4573 (10 nM) for 6 and 24 hours were harvested for cell viability assessment and transcriptomic profiling. (**B**) YX0798 inhibited cell viability in a dose- and time-dependent manner in Mino and JeKo-1 cells. (**C**) The WHITFIELD_CELL_CYCLE_G2_M and HALLMARK_MITOTIC_SPINDLE pathways were significantly downregulated in Mino and JeKo-1 cells upon treatment with YX0798 at 50 and 100 nM. AZD4573 serves as a control. (**D**) Volcano plot shows the significantly altered genes in JeKo-1 cells upon YX0798 treatment at 100 nM for 6 hr. (**E**) Representative genes were significantly downregulated with CDK9i YX0798 or AZD4573. (**F**) mRNA expression of individual genes was validated by quantitative qPCR in Mino and JeKo-1 cells upon treatment with YX0798 and AZD4573 for 6 hr.

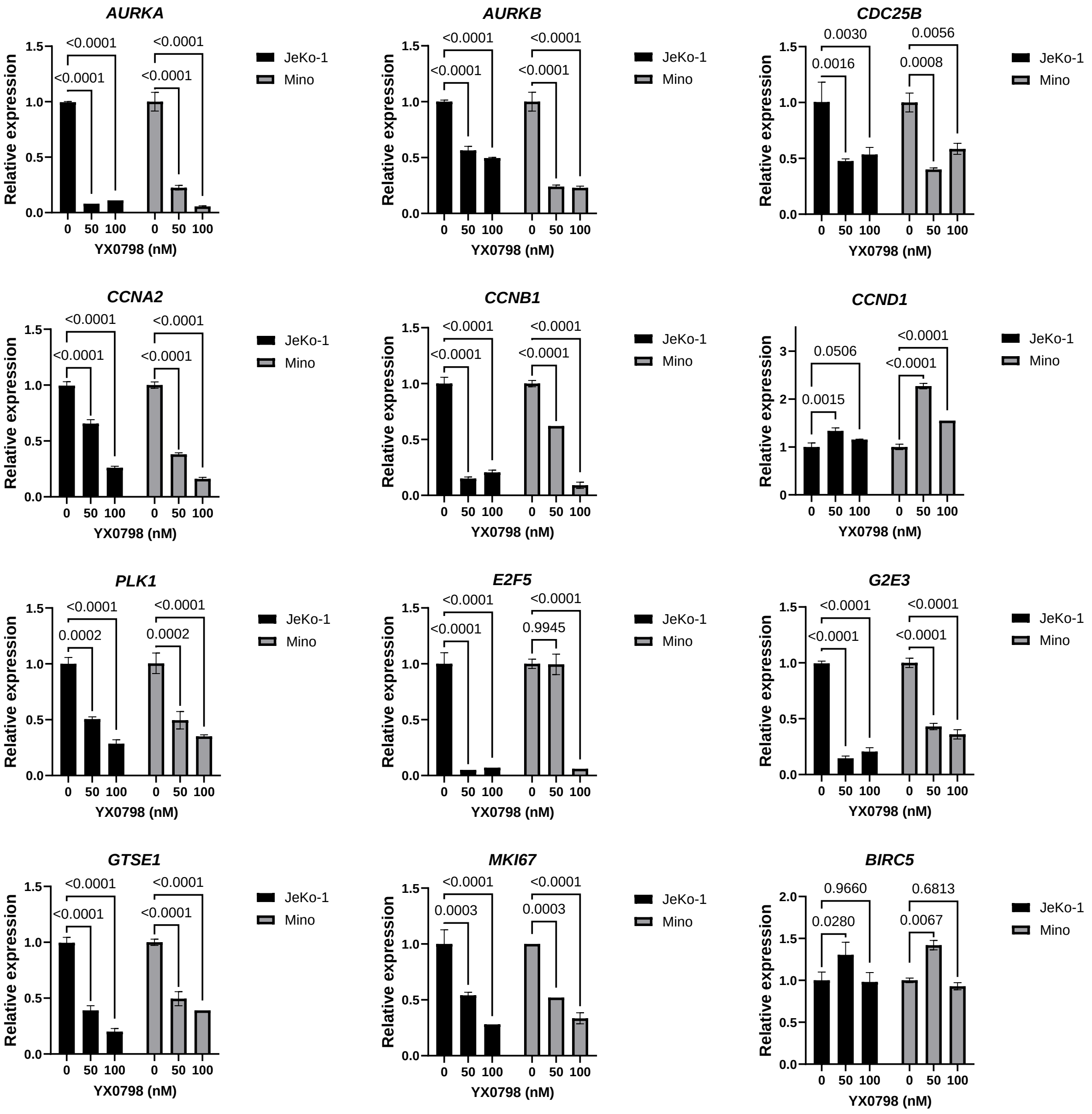


**Figure S3. CDK9i YX0798 downregulates G2/M-specific transcription.**

Quantitative real-time PCR determined the expression of genes indicated in JeKo-1 and Mino cells upon treatment with YX0798 at 50 and 100 nM for 6 hours. Statistical significance (P-value) was generated by comparing YX0798 to DMSO treatment.

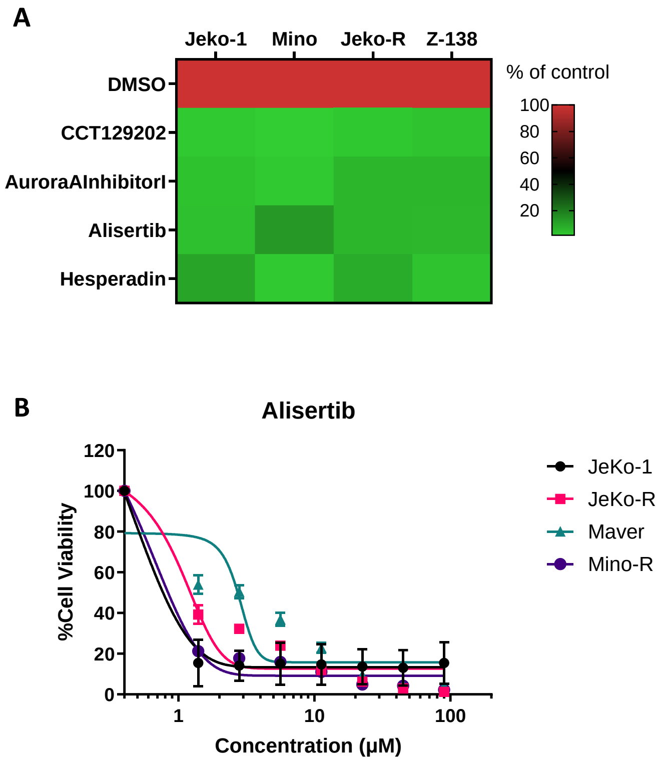


**Figure S4. Inhibition of aurora kinase A is effective in killing MCL cells.**

(**A**) Multiple aurora kinase A inhibitors (n = 4) at 5 μM potently inhibited the cell viability of MCL cell lines (n = 4). (**B**) Aurora kinase A inhibitor alisertib effectively killed MCL cells at 72 hours post treatment.

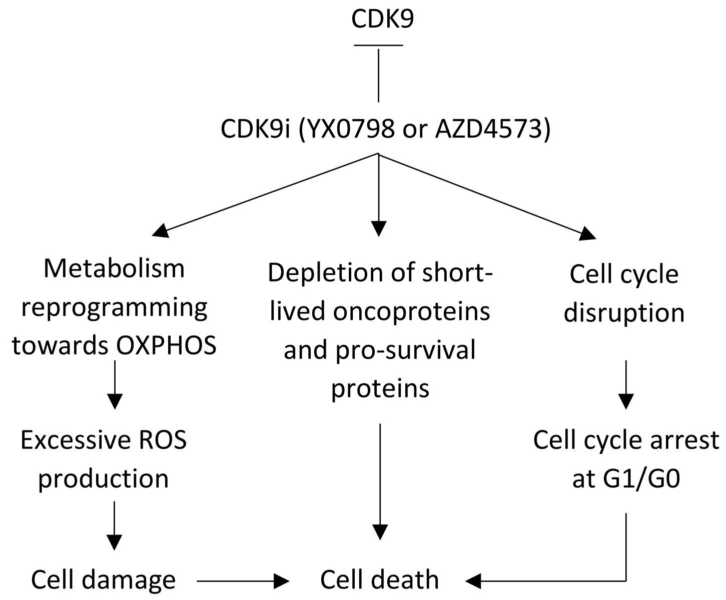


**Figure S5. Proposed models of CDK9 inhibitor YX0798 and AZD4573 in driving MCL cell killing.**
